# Automatic Multi-functional Integration Program (AMFIP) towards All-optical Mechanobiology Interrogation

**DOI:** 10.1101/2021.03.31.437936

**Authors:** Qin Luo, Justin Zhang, Gaoming Lin, Miao Huang, Mai Tanaka, Sharon Lepler, Juan Guan, Dietmar Siemann, Xin Tang

**Author notes:** These authors contributed to the work equally (Qin Luo, Justin Zhang, Gaoming Lin). Corresponding author: Xin Tang, Ph.D.

## Abstract

Automatic operations of multi-functional and time-lapse live-cell imaging are necessary for biomedical studies of active, multi-faceted, and long-term biological phenomena. To achieve automatic control, most existing solutions often require the purchase of extra software programs and hardware that rely on the manufacturers’ own specifications. However, these software programs are usually non-user-programmable and unaffordable for many laboratories. μManager is a widely used open-source software platform for controlling many optoelectronic instruments. Due to limited development since its introduction, μManager lacks compatibility with some of the latest microscopy equipment. To address this unmet need, we have developed a novel software-based automation program, titled Automatic Multi-functional Integration Program (AMFIP), as a new Java-based and hardware-independent plugin for μManager. Without extra hardware, AMFIP enables the functional synchronization of μManager, the Nikon NIS-Elements platform, and other 3^rd^ party software to achieve automatic operations of most commercially available microscopy systems, including but not limited to Nikon. AMFIP provides a user-friendly and programmable graphical user interface (GUI), opening the door to expanding the customizability for many hardware and software. Users can customize AMFIP according to their own specific experimental requirements and hardware environments. To verify AMFIP’s performance, we applied it to elucidate the relationship between cell spreading and spatial-temporal cellular expression of Yes-associated protein (YAP), a mechanosensitive protein that shuttles between cytoplasm and nucleus upon mechanical stimulation, in an epithelial cell line. We found that the ratio of YAP expression in nucleus and cytoplasm decreases as the spreading area of cells increases, suggesting that the accumulation of YAP in the nucleus decreases throughout the cell spreading processes. In summary, AMFIP provides a new open-source and charge-free solution to integrate multiple hardware and software to satisfy the need of automatic imaging operations in the scientific community.

## Introduction

Automatic operations of multi-functional and time-lapse live-cell imaging are essential for biomedical research on dynamic, multi-faceted, and long-term biological questions. Successful automatic operations require streamlined functional coordination of multiple microscope hardware and imaging software systems that are produced by different manufacturers of optoelectronic systems. Most manufacturers usually provide their own hardware-specific drivers and software. Despite their high price, these expensive drivers and software are often non-user-programmable and incompatible with 3^rd^ party hardware, which significantly limits the full utilization of hardware functionalities and the necessary coordination between different devices. To address this unmet need, we have developed a novel software-based automation program, titled Automatic Multi-functional Integration Program (AMFIP), through Java programming language (**Fig. 1**). Compared with other programs available to researchers, AMFIP functions as a hardware-independent controlling hub, enabling the functional coordination of multiple hardware and software systems to achieve automatic multi-functional and time-lapse data acquisition on most commercially available imaging systems. AMFIP is developed based on the μManager platform and entails multiple advantages (**Table 1**).

**Figure 1:**
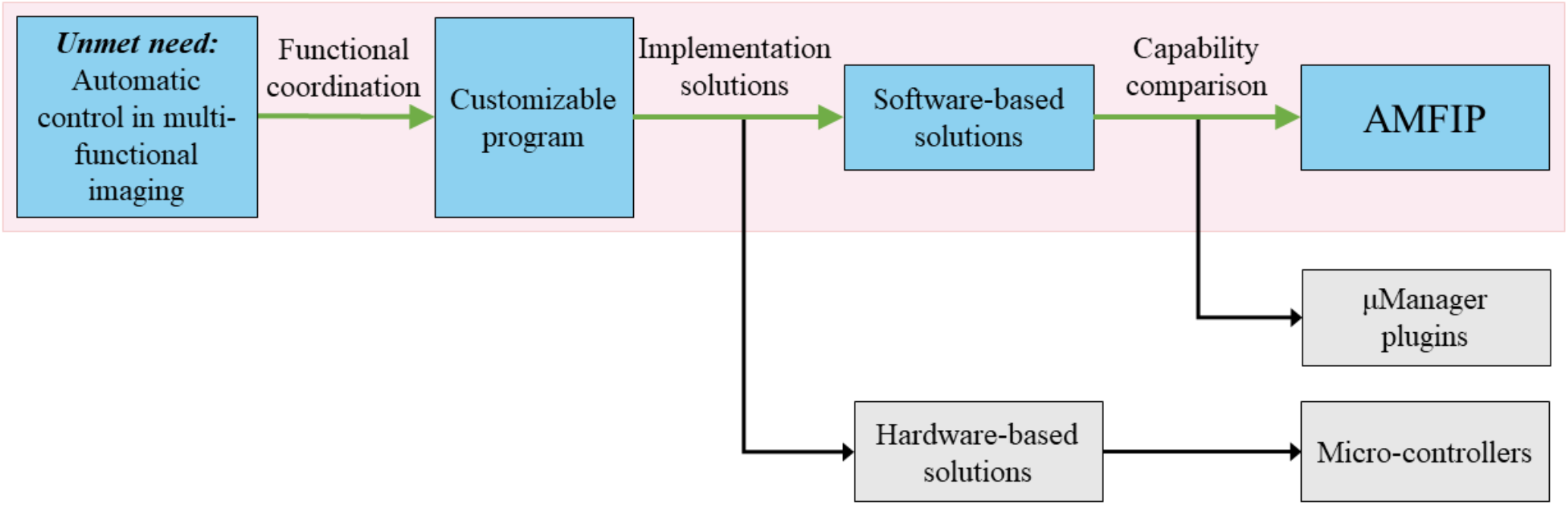
Rationale of AMFIP development. Automatic and customizable control of multiple imaging conditions to acquire spatial-temporal biological information is a capability needed in many research laboratories. In this work, AMFIP, a software-based solution, is developed to achieves customizable and automatic control of generic microscopy systems. In contrast, other software-based solutions, such as μManager plugins, and hardware-based solutions, such as micro-controllers, only control limited hardware and provide restricted functions.

**Table 1:** Comparison between AMFIP and other existing software- and hardware-based solutions. AMFIP coordinates μManager with 3rd party software, such as Elements and SpinView, to manipulate both μManager-supported and non-μManager-supported hardware, such as Nikon A1R confocal microscope system. In contrast, other software-based solutions, such as MultiFRET and EMU, modulate limited μManager-supported hardware, e.g., Nikon TE2000 microscope. AMFIP does not require the purchase of additional hardware. Most hardware-based solutions, such as μSPIM and mmSIM, require NI DAQ card to control

|  | Software-based solutions |  |  | Hardware-based solutions |  |
| --- | --- | --- | --- | --- | --- |
|  | **AMFIP**<br>(Developed in this work) | **MultiFRET**<br>[7] | **EMU**<br>[8] | **μSPIM**<br>[3] | **mmSIM**<br>[1] |
| **Key hardware used for automatic operations** | Nikon Ti2-E inverted microscope (μManager-supported hardware)<br>Blackfly camera (μManager-supported hardware)<br>Nikon A1R confocal microscope system (non-μManager-supported hardware) | Nikon TE2000 microscope (μManager-supported hardware) | Only μManager-supported hardware | Laser from Omicron and Cobolt (μManager-supported hardware)<br>E-665 Piezo Amplifier (μManager-supported hardware)<br>ORCA-flash Camera (non-μManager-supported hardware) | Olympus IX71 microscope (μManager-supported hardware) |
| **Software available for coordination** | μManager, Nikon NIS-Elements, SpinView | Only μManager | | | |
| **Additional hardware to coordinate with optoelectronic devices** | Do not need to purchase any additional hardware |  |  | Need to purchase NI DAQ card (PCI/COM ports required) |  |
| **Achieved automatic functionalities** | Modulation of motor-stage, objectives, filter wheel, light path, and laser<br>Bright-field imaging and confocal microscopy imaging (time-lapse or z-stack)<br>Fluorophore bleaching | High-throughput Förster Resonance Energy Transfer (FRET) image acquisition | High-throughput Förster Resonance Energy Transfer (FRET) image acquisition | Modulation of motor-stage<br>Selective Plane Illumination Microscopy Imaging | Modulation of motor-stage, laser, and filter wheel<br>Structured illumination microscopy (SIM) imaging |

First, AMFIP achieves software-based automation without adding any extra hardware. To achieve automatic operations, some existing solutions require extra single-board microcontrollers to coordinate multiple hardware and software systems. These microcontrollers generate analog and digital signals to modulate optoelectronic devices. For example, the NI data acquisition card (DAQ; National Instrument Corp.) is utilized to control the frame acquisition of the camera and switch of the laser channels.[1] An Arduino-based system is applied to modulate selective-plane illumination microscopy (SPIM).[2] However, these hardware-based solutions need additional purchases of expensive adaptive drivers at prices around the $1000s for some optoelectronic devices, such as the Nikon A1R controller. Our hardware-independent AMFIP program avoids such additional expenses by implementing a home-built Java-based script that coordinates all devices through software communications alone. Traditionally, to transmit modulatory signals from the NI digital-analog converters (DAC) card into some optoelectronics hardware, particular physical ports are needed, such as Communication (COM) port or Peripheral Component Interconnect (PCI) port. However, these ports are not always present in many commercial devices, such as the Nikon LU-N4 laser units used by many research laboratories. Thus, additional purchases are needed for researchers to control the new hardware components that contain these ports.[3] AMFIP bypasses these hardware constraints and accomplishes automatic modulations by leveraging the software communications between the Nikon NIS-Elements software platform (Element) that exclusively controls Nikon’s hardware and the hardware equipment from 3^rd^ party manufacturers.

Second, compared to other existing solutions, AMFIP supports a wider range of hardware, including but not limited to non-μManager-supported hardware. As an open-source software package, μManager has been applied to manipulate optoelectronic devices.[4]–[6] Researchers have developed various user-defined μManager plugins to achieve specific tasks. However, due to the rapid upgrades of equipment in the market, most existing μManager plugins lack speedy enough development to provide sufficient compatibility with the latest microscopy equipment. For example, a recently developed μManager plugin, MultiFRET, can achieve automatic acquisition and analysis of fluorescent images, but relies on μManager-supported instruments.[7] Due to the same restriction, another newly developed plugin, Easier Micro-Manager User (EMU), can offer only limited functions to control μManager-supported optical hardware, e.g., modulation of laser and filter wheels, and acquisition processes, e.g., time series or z-stack imaging.[8] In contrast, our AMFIP can work with non-μManager-supported instruments, providing more choices on new hardware to be included in microscope systems. We have demonstrated that AMFIP enables automatic operations such as modulations of the Nikon laser channels and multi-functional imaging (**Results (F)**), which have been previously unattainable by the μManager platform. [4]

Third, Java-based AMFIP enables user-programmable and automatic operations. Specifically, AMFIP has a user-friendly GUI and is programmable to perform operations that meet the experiment-specific requirements. Prior to experiments, researchers can flexibly specify the functions needed for their experiments and input the required parameters. Next, AMFIP coordinates all hardware and software involved in the experiments and executes the specified functions automatically. In case unexpected accidents occur during the course of experiments, researchers can flexibly pause functions without any loss of acquired data and resume the experiments at any time after clearance of the accidents. For comparison, an existing MATLAB-based GUI[9] can automatically accomplish an 18-hour data acquisition, but it cannot pause the experiments in case of unexpected accidents. Instead, AMFIP achieves automation of the entire experiment while providing users with safe and flexible control. These features are desired for both new and experienced users.

In summary, our AMFIP program provides a charge-free and hardware-independent solution for multi-functional and long-term image acquisition in biomedical research. To achieve automatic control of multiple optoelectronic hardware, AMFIP coordinates μManager with the Elements platform and other 3^rd^ party software. Its user-friendly GUI allows researchers to flexibly customize and program AMFIP to meet different experimental requirements.

## Results

### A. Graphical User Interface (GUI) of AMFIP

A user-friendly and easily understandable program will benefit new users to effectively start their research activities. Based on the Application Program Interface (API) of μManager that prompts users with all inputs of experimental parameters and essential functions available, AMFIP presents such an all-in-one graphical user interface (**Fig. 2**) to prescribe and implement multi-functional data acquisition, such as coordination of multiple field-of-views (FOVs), selections of microscope objectives, and modulation of multiple laser channels. Simultaneously, AMFIP preserves full access to other 3rd party software and allows real-time adjustments to achieve customized configurations.

**Figure 2:**
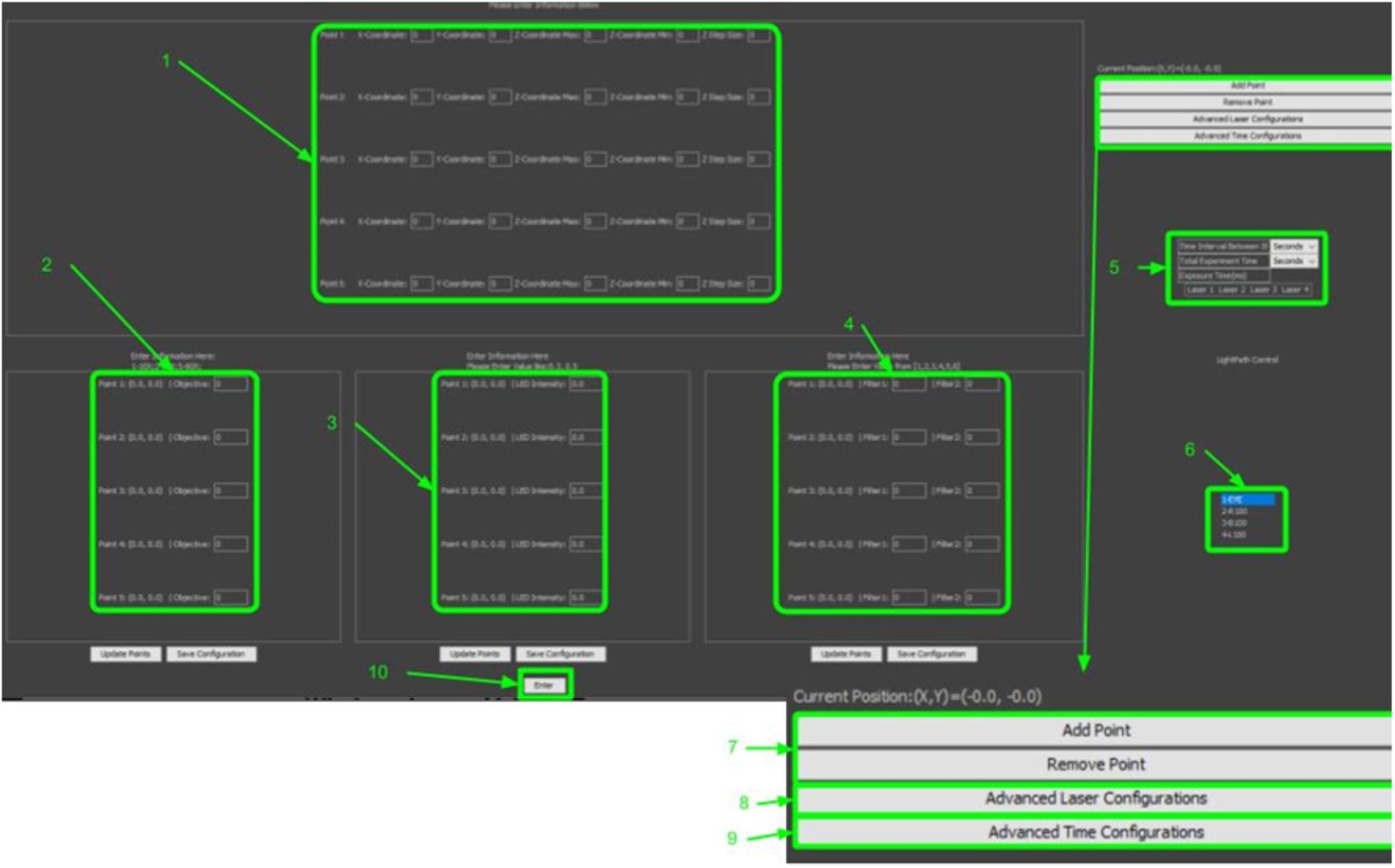
The Diagram of the GUI of AMFIP. **(1)** Input X, Y, Z coordinates for each FOV. **(2)** Specify objective type for each FOV. **(3)** Specify DiaLamp intensity for each FOV (Used for BF images). **(4)** Specify desired filter cubes for each FOV. **(5)** Universal time and laser configurations for each FOV in the experiment. **(6)** Light Path Control. **(7)** Buttons to add or remove points. **(8)** Button to open up the laser configurations window. **(9)** Button to open up additional time configurations window. **(10)** Enter button to signal the start of the specified experimental procedure.

### B. Design Rationale and Structure of AMFIP

The workflow of AMFIP follows a model-GUI-controller paradigm that consists of 3 compartmentalized and interconnected logical components: the model (data), the GUI, and the program controller (**Fig. 3**). This paradigm of compartmentalized components enables smooth implementation of new functions by allowing the users to customize any component(s) without interfering with others. For example, users can flexibly change the GUI of AMFIP to a new layout without modifying the model or program controller.

**Figure 3:**
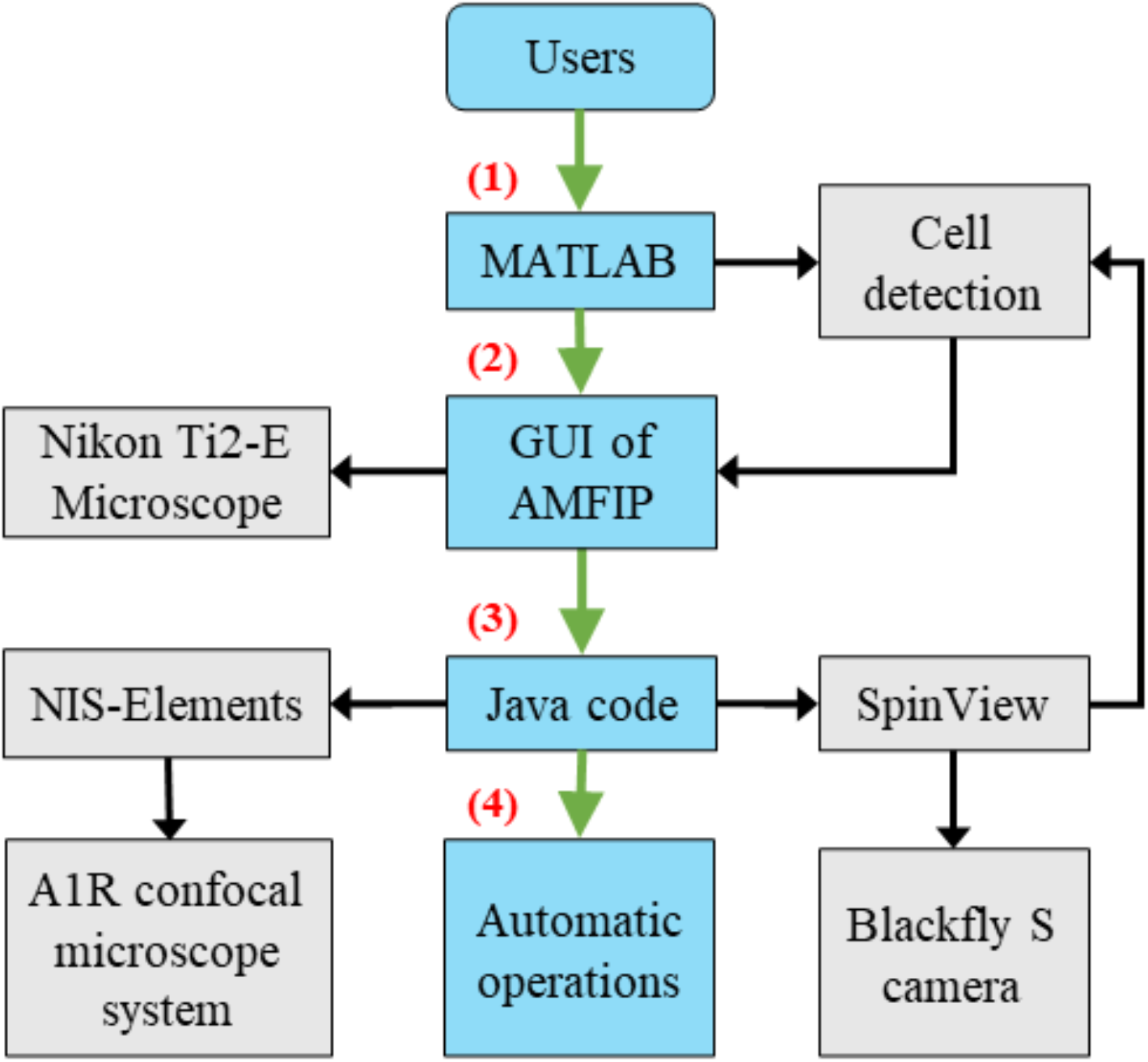
The Workflow of implementing AMFIP in a typical experiment. **(1)** A MATLAB program detects the cells of interest and generates a coordinate list of FOVs. **(2)** The GUI of AMFIP reads the coordinates to guide the movement of the XY motor-stage. Users input predefined experimental parameters and initiate μManager to modulate the Nikon Ti2-E microscope. **(3)** Java code in AMFIP activates Elements and SpinView to manipulate the A1R confocal microscope system and Blackfly S camera. The MATLAB program reads the latest bright-field image to update the coordinates of current FOVs. **(4)** AMFIP automatically conducts these operations in sequence for multi-functional biological imaging.

At the beginning of experiments, users input experimental parameters into the GUI. Once finished, the controller retrieves the data from the GUI, updates the model, and saves the data to local hard disk. Alternatively, users can select a previously saved .JSON configuration file. This selection will restore a previously saved configuration into the input fields of the GUI, but will not execute them right away. This feature gives the user the full control to adjust any values of experimental parameters if needed. Alternatively, if a user is entering a new configuration, she/he can choose to save the current configuration for future use. This feature is designed to enable robust repeatability for experiments that take place at different time points and allow other users to replicate and corroborate experiments. Next, the program controller retrieves the inputted data from the model and conducts the data-specified experiments accordingly.

For complex experiments where users want to frequently pause the experimental procedure to adjust for new conditions/samples, such as adding a pharmacological drug into the cells, AMFIP allows users to inform the hardware when to pause and resume in a user-determined manner (**Fig. 2**). Once manipulation of the conditions/samples is done, users can restore the experiment progress by clicking the “resume” button on the GUI (**Fig. 2**). The program will automatically pick up where it left off and continue the pre-defined procedure. AMFIP also has a manual pause function incorporated that can be used by users to pause whenever they desire. This feature may serve as an emergency stop.

### C. Functional Synchronization of Nikon NIS-Elements and μManager

Nikon NIS-Elements platform is a single and universal software that exclusively controls Nikon’s hardware. Currently, it is the only commercially available software that controls the Nikon A1R confocal microscope system for 3D image acquisition. However, utilizing its full automatic imaging functionalities requires purchases of additional Nikon hardware and software. Additionally, the built-in macro inside Elements only achieves automatic image acquisition with limited functionalities.[10] To overcome this restriction, we designed the userfriendly GUI of AMFIP so that users can directly input and compile a sequence of macro commands into a single text field. A macro consists of a sequence of executable commands that utilize a set of predefined functions in Elements, e.g., the switch of laser channels, adjustment of laser intensity, or acquisition of 3D z-stack images. This text field generates a *.mac file that is saved in a specified directory in the local computer. This *.mac file can be loaded into Elements to execute predefined functions of the hardware. However, a macro only controls the internal functions of Elements and is incompatible with non-Nikon hardware and software.

To overcome this limitation and to achieve automatic operations, AMFIP enables the functional synchronization of macros and μManager (**Fig. 3**). To use the program, we first set a configuration that defines an automatic sequence of motor-stage movements through the GUI. For each FOV to which the motor-stage moves, a script in AMFIP will activate Elements and run a series of commands in the macro editor to execute the experiment-specific functions of the Nikon A1R confocal microscope, such as laser illumination and fluorescent imaging. Upon the completion of image acquisition at each FOV, the macro will generate a blank report and save it into a user-defined file directory. Meanwhile, AMFIP continues checking for this blank report file. Once the program finds this report file, the motor-stage moves to the next FOV and the program deletes the previous file. This process will be automatically repeated in a programmable manner until all pre-selected FOVs are imaged and saved.

### D. Race-hazard-free Coordination between SpinView and μManager

AMFIP can utilize the Blackfly camera that is controlled by SpinView, a GUI provided by FLIR©, to conduct bright-field image acquisition. However, since SpinView cannot directly communicate with other 3^rd^ party software without additional programming, a race hazard between imaging and stage movement may occur during experiment processes. To avoid this potential hazard and achieve safe operations, AMFIP connects SpinView and μManager via Java codes.

Specifically, AMFIP contains a home-built script that controls the keyboard and the cursor of PC to manipulate SpinView (**Fig. 3**). After μManager executes several automatic operations, such as motor-stage movements and the switch of microscope objectives, the Java code launches SpinView automatically to acquire and save a bright-field image within 1 second. To avoid the condition that SpinView-regulated image acquisition is perturbed by μManager-regulated stage movement, we pre-allocate a waiting time (usually ~ 2 seconds) during which μManager pauses its operations. This pre-allocated waiting time is sufficient for the operations of SpinView and does not overtly extend the duration of the total experiment. After each image has been captured and saved, μManager resumes its own predefined tasks and repeats this cycle till the completion of the experiment. AMFIP is flexible in adding user-compiled scripts either before or after the start of macro in Elements, allowing different experiment requirements to be met in a crosstalk-free manner.

### E. Connections between MATLAB and μManager for Automatic Cell Detection

AMFIP coordinates with MATLAB to realize precise detection of cell samples and timely prevention of motor-stage drift during experiments (**Fig. 4**). The first function aims to find all FOVs that contain cells of interest and guide the XY-plane movement of the motor stage to these spots. The second function is to monitor the current coordinates of FOV and to ensure that the cells of interest are captured even if their positions drift during experiments. To achieve frequent and real-time communications between μManager and MATLAB, AMFIP connects the two software by implementing its Java codes.

**Figure.4:**
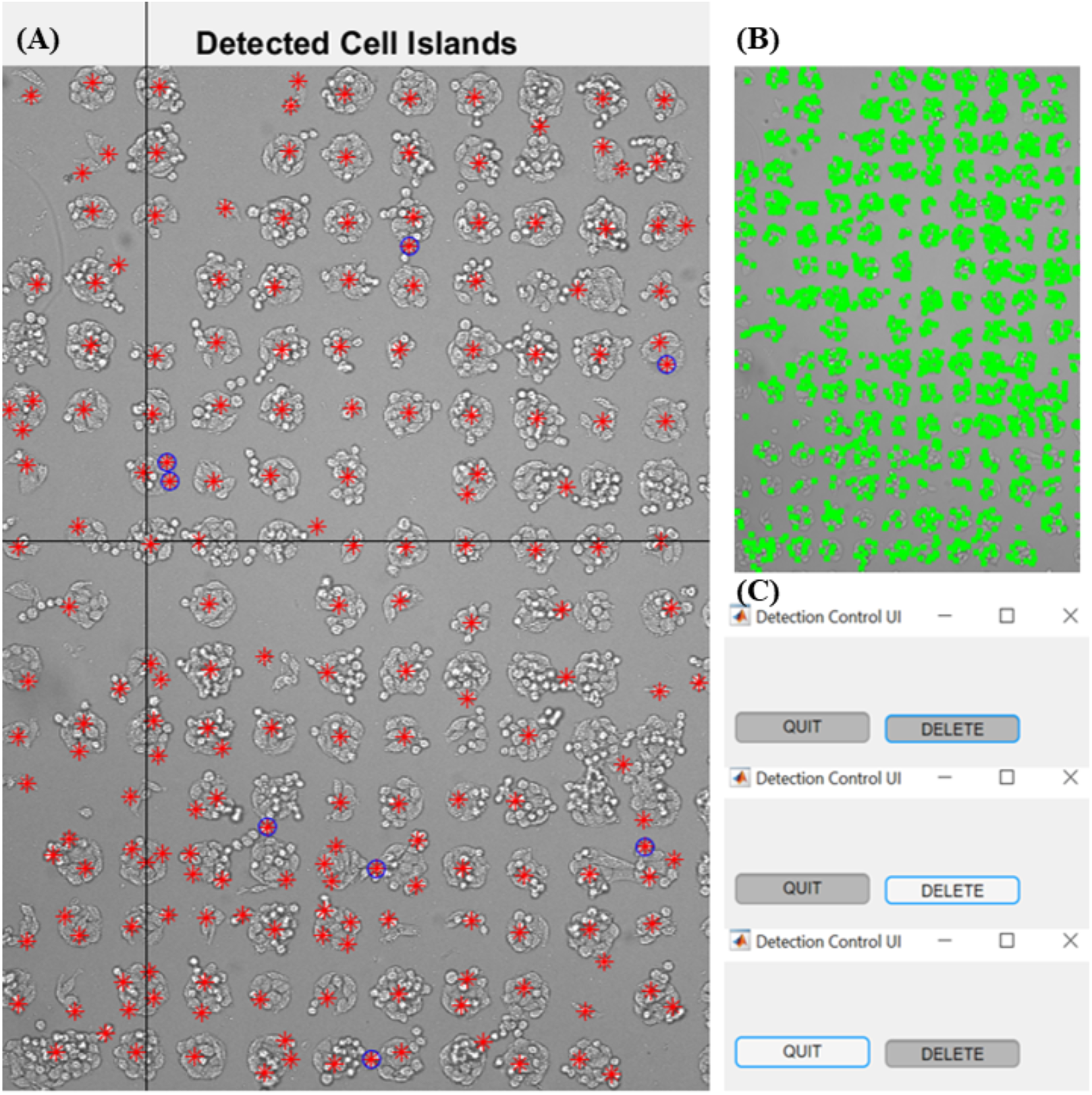
A representative view of MATLAB operation interface for automatic cell detection and stage-drift prevention. **(A)** Interface to edit marks of detected cell islands. Red marks are the centroids of detected cell islands while blue circles are unwanted marks **(B)** Pattern of detected cells. The detected cells are marked with green rectangles. **(C)** Detection Control UI with “DELETE” and “QUIT” buttons. The grey button means that its statement is inactivated. Click the button and its function is activated.

Similar to the control of SpinView, AMFIP utilizes Java code to control the keyboard and the cursor of PC to operate on the interface of MATLAB. For precise detection of cell samples, we designed four steps:

1. Use a Blackfly S BFS-U3-70S7M camera to take a bright-field image at 10× magnification under the control of μManager.
2. Launch the MATLAB program by AMFIP to read the image and to distinguish cells from non-cell matters by image processing, such as dilating, smoothing, edge detection, and segmentation.
3. Edit the image, remove incorrect markers on non-cell matters, and add new markers on detection-missed cells on the user-friendly MATLAB GUI.
4. Analyze the edited image by the MATLAB program to generate a text-format list of the coordinates of the selected markers that locate on the cells’ centroid. This text-format list can be read by μManager to guide the movement of the motor-stage on the XY-plane.

For the second function, i.e., monitoring the coordinates in situ, the MATLAB program maintains a specified directory to temporarily save and transfer all captured images one by one. At any given moment, there is only one image file present in this directory. AMFIP constantly monitors this directory, reads this image, and transfers this image to a destination folder. During this process, the MATLAB directory constantly contains only the latest image that is used to accurately update the current coordinates of FOV. This monitoring function is combined with the AutoFocus functions (Z-axis) provided by the Elements to ensure precise 3D time-lapse imaging for long-term biomedical experiments.

### F. Multi-functional and long-term imaging of Beas2B (B2B) cell line that expresses Yes-associated-protein (YAP)

To verify the capability of AMFIP in practical experiments, we designed and conducted a series of multi-channel and long-term image acquisition experiments to elucidate the dynamics of cell spreading - a fundamental mechanobiological phenomenon. Specifically, in this experiment, we aimed to co-track the spatial-temporal dynamics of YAP expression in cells and the traction force applied by cells throughout the process of cell spreading on a 2D hydrogel substrate. As a mechano-sensitive protein in cells, YAP, along with transcriptional coactivator with PDZ-binding motif (TAZ), shuttles between the nucleus and cytoplasm depending on the specific mechanical signals received, such as the cell traction and microenvironment stiffness.[11] However, the quantitative relationship between the ratio of YAP expression in the nucleus/cytoplasm and cell traction during cell spreading remains unknown, largely due to the lack of tools that enables simultaneous recording of real-time YAP expression and cell traction. In this work, we combined AMFIP, YAP-expression human bronchial epithelial B2B cells [12], and traction force microscopy to fill this knowledge gap.

We found that, during the spreading process of single B2B cells on the 2 kPa hydrogel substrate, YAP expression in the nucleus decreases in comparison to that in the cytoplasm (n = 5; **Figs. 5A & 5C)**. For B2B cells that flattened down from suspension state to adherent state during the first 9 hours of experiment, the average normalized ratio of YAP nucleus/cytoplasm intensity (N/C ratio) changed from 1 to 0.76 ± 0.045 (n=5; p-value = 0.0002***; **Figs. 5C1 & 5C2**), while the average normalized cell spread area steadily increased from 1 to 1.81 ± 0.141 (p-value < 0.0001****; **Fig. 5C1**) and the average normalized nucleus spread area simultaneously augmented from 1 to 2.00 ± 0.136 (p-value = 0.0079**; **Fig. 5C2**). The results suggest that YAP in the nucleus of single B2B cells may translocate into the cytoplasm as the spread areas of both the cell body and the nucleus increase throughout the cell spreading process.

**Figure 5:**
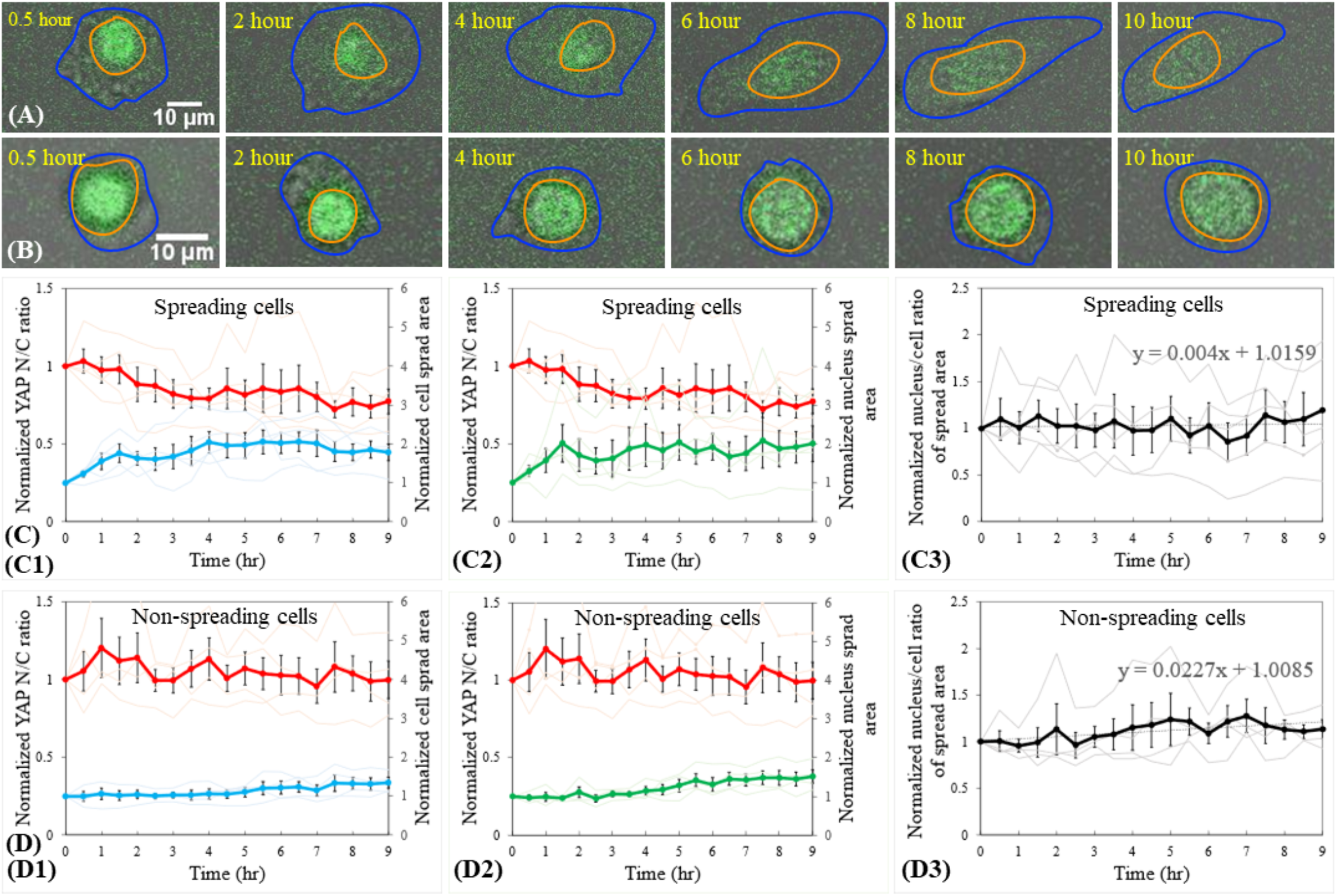
Relationship between YAP N/C ratio and cell spreading states of single B2B cells. **(A)** 10-hr time-lapse image stack that contains overlapped fluorescent YAP images and bright-field cell images of a single spreading cell. Note: The shapes of the cell (in blue contour) and nucleus (in red contour) transform from roundness to flatness. Normalized YAP N/C ratio decreases from 1 at 0th hour to 0.56 at 10th hour. **(B)** 10-hr time-lapse image stack of a nonspreading single cell. Note: The shape of the cell maintains rounded. Normalized YAP N/C ratio changes from 1 at 0th hour to 1.04 at 10th hour. **(C)** YAP N/C ratio versus nucleus/cell-body area of spreading cells (n=5). The average normalized YAP N/C ratio (red bold line; n=5) changes from 1 to 0.76 ± 0.045 (p-value = 0.0002***; **C1**). In parallel, the average normalized cell area (blue bold line; n = 5) increases from 1 to 1.81 ± 0.141 (p-value < 0.0001****; **C1**) and the average normalized nucleus area (green bold line) changes from 1 to 2.00 ± 0.136 (p-value = 0.0079*; **C2**). The average normalized nucleus-projected-area/cell-projected-area ratio approximately remains constant at 1 ± 0.043 (p-value = 0.6398 (ns (not significant))), with a trend-line slope of 0.004 **(C3)**. **(D)** YAP N/C ratio versus nucleus/cell-body area of nonspreading cells (n=4). The average normalized YAP N/C ratio (red bold line) practically stands at 1 ± 0.031 (p-value = 0.7422 (ns); **D1**), and the average normalized cell area (blue bold line) increases from 1 to 1.35 ± 0.064 (p-value =0.0113*; **D1**). The average normalized nucleus area (green bold line) rises from 1 to 1.51 ± 0.106 (p-value = 0.0010***; **D2**) The average normalized nucleus-projected-area/cell-projected-area ratio changes from 1 to 1.13 ± 0.048 (p-value = 0.1519 (ns)), with a trend-line slope of 0.0227 **(D3)**.

To validate that the changes in the YAP N/C ratio are independent of the changes in cell morphology, we calculated the normalized ratio of nucleus-projected-area/cell-projected-area during cell spreading. We found that this ratio approximately changed from 1 to 1.19 ± 0.043 over time (p-value = 0.6398 (ns (not significant)); **Fig. 5C3**), with a trend-line slope of 0.0004. The results suggest that the sizes of both nucleus and cell body may increase proportionally. Based on this evidence, we reason that the decline of YAP N/C ratio may not be a result of the disproportionality between the changes of nucleus area and cell spread area. We suppose that the decrease of the YAP N/C ratio in single B2B cells throughout the cell spreading process may be caused by the translocation of YAP from the nucleus to the cytoplasm.

As the control, we examined the single B2B cells that did not spread (n=4) in the same experiments. We found that the average normalized YAP N/C ratio nearly maintained a constant value at 1 ± 0.031 (p-value = 0.7422 (ns); **Figs. 5D1 & 5D2**), while the average normalized cell spread area increased from 1 to 1.35 ± 0.064 (p-value =0.0113*; **Fig. 5D1**) and the average normalized nucleus spread area rose from 1 to 1.51 ± 0.106 (p-value = 0.0010***; **Fig. 5D2**). These results suggest that non-spreading single B2B cells expand by a smaller degree (43% of the increase) in size compared with spreading cells (91% of the increase), while their ratio of nucleus-projected-area/cell-projected-area almost remains stable. For nonspreading single cells, the average normalized nucleus-projected-area/cell-projected-area ratio increased from 1 to 1.13 ± 0.048 (p-value = 0.1519 (ns); **Fig. 5D3**), with a trend-line slope of 0.0227. We reason that fewer YAP, if any, shuttles from nucleus to cytoplasm in non-spreading single B2B cells in comparison to spreading single cells throughout the adhering-to-spreading processes.

Together, the results suggest that the YAP N/C ratio in B2B single cells may depend on the spreading state of cells - after 8-9 hours of experiment, the average normalized YAP N/C ratio is 0.74-0.76 in spreading cells and is 0.99-1.00 in non-spreading cells (p-value = 0.0113*; **Figs. 5C1 & 5D1**). These mechanobiological results suggest that a nucleus-to-cytoplasm translocation of YAP may occur in response to the spreading states of B2B single cells (**Fig. 5**). In this experiment, AMFIP demonstrates the verified multi-functionality and stability for data acquisition that require minimum manual operations.

## Discussion

In this work, we introduce a verified automation program (AMFIP) to overcome the limitations of the latest solutions for image acquisition in biomedical research (**Table 1**). Currently, most manufacturers of optoelectronic hardware provide their own software with limited charge-free opportunities for functional customization and automation.[13] These software programs exclusively control manufacturers-specified hardware and require additional financial expenses to coordinate with different microscopy systems.[13], [14] In contrast, AMFIP functions as a control hub that coordinates multiple software programs to enable modulation of different microscopy systems. Taking our experiments described above as an example, AMFIP can automatically integrate and execute the following functions in a user-defined sequence (**Fig. 3**). First, AMFIP communicates with μManager to modulate the movement of the XY motor-stage. Second, the Java code of AMFIP activates SpinView to acquire and save a bright-field image. Third, the MATLAB program embedded in AMFIP automatically generates a list of coordinates of FOVs based on the acquired bright-field image. Finally, AMFIP reads this list and guides the movement of the XY motor stage based on the coordinates that are constantly updated by the cell detection program.

Further, the execution of specified Java code embedded in AMFIP functions as a bridge connects the user’s inputs into AMFIP with the functional and automatic implementation of 3rd party software. Therefore, once users develop the Java-code-enabled coordination between AMFIP and other 3^rd^ party software, these coordination activate the existing automatic functions inside these 3^rd^ party software. As a result, users can leverage the existing customizability and automation inside many commercial software programs, such as a macro in Elements, and do not need to compile additional home-built functions starting from scratch, which is either time-consuming or technically challenging.

In addition to verifying the performance of AMFIP, we chose co-tracking the YAP N/C ratio and cell traction as an example because mechanistic elucidation of the mechanosensitive YAP/TAZ will deepen our understanding of tissue development, healthy homeostasis, and cancer progression. [15], [16] The nuclear accumulation of YAP/TAZ dominates their active interactions with transcription factors, such as TEADs, RUNXs, and p73, to regulate the transcription of specific genes that influence cell proliferation, migration, and survival.[15], [17] In particular, emerging evidence suggests that YAP/TAZ show oncogenic effects in most cancer types and tumor-suppressive effects in some cases. In the former case, high protein and mRNA levels of YAP/TAZ are found to be associated with the poor prognosis of cancer patients.[18]–[22] As the mechanical microenvironment of tumors is dramatically different from that of healthy tissues[23]–[31], we suppose that mechanosensitive YAP/TAZ may be aberrantly expressed in the neoplasia niche and activate the oncogenes to promote malignant transformation. Our AMFIP enables visualization of the real-time interplay (a “dance”) between YAP N/C ratio and cell mechanics and provides one step further towards understanding how YAP/TAZ mediates tumor biology. These AMFIP-enabled *in vitro* results may inform the future directions of *in vivo* research and ultimately guide the innovation of new cancer therapies.

## Conclusion

In summary, we have developed AMFIP, a hardware-independent and software-based program for automatic data acquisition that is applicable to many microscopy systems and optoelectronic hardware. We have verified the capability of AMFIP by demonstrating the relationship between YAP dynamics and cell spreading states in a human epithelial cell line. We believe AMFIP is a reliable platform that could benefit the scientific community. The source code of AMFIP is charge-free and available to the public on GitHub (link: http://). We hope users will develop and expand more new functions based on AMFIP to enhance their specific automatic operations in research.

## Materials and Methods

### 1. Hardware and Software

Hardware systems used for experiments include a commercial fluorescent confocal microscope system (Nikon A1R HD25), a monochrome camera (BFS-U3-70S7M-C FLIR©), and a desktop computer that is installed with a 64-bit Microsoft© Windows 10 Pro operating system. The Nikon confocal microscope system consists of multiple components: Ti2-E inverted microscope, LU-N4 laser unit (405 nm, 488 nm, 561 nm, and 640 nm laser channels), confocal controller, standard fluorescence detector (4 photomultiplier tubes (PMT) and 6 filter cubes), and a scan head (2 galvano scanners and 1 resonant scanner). AMFIP controls the confocal imaging components, such as the laser unit, confocal controller, detectors, and scan head, through activating the Elements. Ti2-E inverted microscope comprises a LED Lamp-house for illumination, motorized XY stage, 6 motorized epi-fluorescence filter turrets, 7 motorized condenser turret, 6 motorized nosepieces, and a Stage joystick. AMFIP controls the Ti2-E inverted microscope through coordinating with μManager.

Confocal 3D image stacks and videos are acquired by the confocal microscope system. The Ti2-E inverted microscope works independently to the acquisition of bright-field images by the monochrome camera. The Dialamp (a white LED equipped on the Ti2-E microscope) serves as a light source for bright-field imaging.

Three software systems are involved to automatically coordinate these devices: (1) SpinView, which controls the BFS-U3-70S7M-C camera; (2) Nikon NIS-Elements, which controls the whole confocal system; and (3) μManager, a 3rd party software, which controls the independent operation of Ti2-E microscope. AMFIP controls all these three software systems through the IntelliJ IDEA platform.

### 2. AMFIP Guideline

#### 2.1 Setting up the programming environment

The following steps show how to set up the software environment to program and implement AMFIP:

**2.1.1** Download and install μManager software from https://micro-manager.org/wiki/Download%20Micro-Manager_Latest%20Release. The latest version μManager 2.0-gamma is recommended because of its active development and maintenance.
**2.1.2** To coordinate μManager with optoelectronic hardware: (1) Connect all needed optoelectronic hardware to a desktop computer and turn on these hardware systems. (2) Add the adaptive drivers called “device adaptor” of the optoelectronic hardware provided by either μManager or the hardware manufacturer into the μManager directory; (3) Go to “Devices->Hardware Configuration Wizard”, check “Create new configuration”, and click “Next”; (4) Find the names of connected hardware in “Available Devices”, click “Add”; (5) A confirmation window pops up. Check their properties and click “OK”; (6) A “Peripheral Devices Setup” window pops up. Select all needed peripheral devices of parent devices(the connected hardware) and click “OK”. These peripheral devices are configured in the list of “Installed Devices”. Click “Next”; (7) Select the default devices and click “Next”; (8) (optional) Set delays for devices without synchronization capabilities and click “Next”; (9) (optional) Define position labels for state devices, such as filters and objectives, and click “Next”; (10) Save the new configuration file, restart μManager, select this configuration file in “Micro-Manager Startup Configuration” and click “OK”.
**2.1.3** Enable the control of all connected and configured optoelectronic hardware by μManager. For example, we control the Nikon Ti2-E microscope by μManager in our lab. The adaptive driver of Ti2-E microscope is “Ti2_Mic_Driver.dll ‘’ located in the “Nikon\Ti2-SDK\bin” of the Ti2 Control software’s directory. This software can be downloaded from https://www.nikon.com/products/microscopesolutions/support/download/software/biological/. Ti2 Control Ver 1.2.0 rather than the latest version is recommended because of its better compatibility with μManager in Microsoft Windows 10 operating system.
**2.1.4** Download and install IntelliJ IDEA from https://www.jetbrains.com/idea/download/#section=windows for the development of Java-based software.
**2.1.5** Download and install Java Development Kit (JDK) from https://www.oracle.com/java/technologies/javase-jdk15-downloads.html. JDK 14.0 or higher version is recommended for programming AMFIP.
**2.1.6** Set up software configuration in IntelliJ to allow developing μManager-based programs. First, open IntelliJ and go to “Settings->Compiler->Annotation Processors”. Check the box of “Enable annotation processing”. Second, go to “Project Structure- >Artifacts” and create a JAR(Empty) file. The output directory should be the mmplugins folder on the μManager directory. Third, go to “Project Structure- >Libraries”, add “mmplugin” and “plugins/Micro Manager” folder from the directory of μManager.
**2.1.7** Click “add Configuration” and create an application with the following information: “ Main class: ij.ImageJ; VM option: -Xmx3000m -Dforce.annotation.index=true; Work directory: μManager directory; Use classpath of module: the name of current project”.
**2.1.8** Click “Run” in IntelliJ to launch μManager. Click “OK” and the main interface of μManager appears.

#### 2.2 Use of GUI (Fig. 2)

The following steps show how to input pre-defined experimental parameters into the GUI of AMFIP and start a multi-task experiment.

**2.2.1** Open μManager, the GUI of AMFIP is under “Plugins->Automation”.
**2.2.2** Define the number of FOVs to which XY motor-stage moves by clicking “Add Point” or “Remove Point”. Input the coordinates of each FOV into text fields under “Coordinate Panel”. Alternatively, retrieve the saved configurations, i.e., JSON files with a list of previous experimental or pre-defined parameters, including the number/coordinates of FOVs, imaging conditions, and data acquisition parameters.
**2.2.3** Input a quantitative value into the “Total Experiment Time” text field to define the entire duration of the experiment. Click “Additional Time Configurations”, and input quantitative values into “Start Time”, “Time Interval” and “End Time” for each specified FOV. For each FOV, click “Pause” to program the time when the experiments should automatically stop. The experiments can be resumed by manually clicking “Resume”.
**2.2.4** Modulate microscope objectives, DiaLamp, and excitation/emission filters for each FOV by inputting predefined quantitative values into the three sub-panels below “Coordinate Panel”.
**2.2.5** Next, click “Save Configuration” to keep a record. Under the submenu of “Devices”, the window of “Device Property Browser” presents a list of quantitative values as a reference, e.g., value “1” for the configuration of microscope objective refers to switching to 10× objective for current FOV.
**2.2.6** Once all parameters are fed into the GUI, click “Enter” to start a task.
**2.2.7** In case some unexpected conditions occur during the experimental process, click “Pause” to temporarily stop the experiment. The experiment can be resumed by clicking “Resume”.

#### 2.3 Use of home-built Java code to coordinate μManager, NIS-Elements, and SpinView

The following steps show how Java code is applied to automatically coordinate μManager with other software.

**2.3.1** Open the AMFIP’s Java project in IntelliJ and go to “src”. Scripts are created in “CameraScript” and “ElementsScript” .java files.
**2.3.2** Go to “Main->Executor”, add two statements: “CameraScript.main()” and “ElementsScript.main()” into “scheduleTaskForAPoint” function. “ElementsScript.main()” activates Elements and runs a predefined macro. “CameraScript.main()” activates SpinView to automatically capture and save the bright-field images.
**2.3.3** To control NIS-Elements by Java code, maximize the window of Elements to enclose the window of AMFIP GUI, and enable cursor-based activation of Elements functions. To activate SpinView, place the icon of this software into the taskbar and control the camera by Java code.
**2.3.4** Initiate part 2.2.5. Once XY motor-stage moves, Elements and SpinView are launched and will enable automatic hardware operations following the predefined commands in Java code.

#### 2.4 Use of MATLAB program for cell detection

Automatic cell detection is achieved by our MATLAB program embedded in AMFIP. The following steps show how our MATLAB program cooperates with AMFIP to achieve cell detection function during the experiment.

**2.4.1** Our MATLAB program is integrated into the AMFIP package (GitHub: Http://). Unzip this package, move the MATLAB program to the specified working folder for bright-field images, and add this folder to the MATLAB path.
**2.4.2** Create a text file named “data.txt” in this folder.
**2.4.3** Run the installed AMFIP to start the experiment in IntelliJ IDEA. In the GUI of AMFIP, the user sets up the original FOV, where the first 10× bright-field image is taken. After the camera finishes taking the first 10× bright-field image, our MATLAB program will be activated by AMFIP.
**2.4.4** A figure window of the first 10× bright-field image and a MATLAB GUI pop up. On the figure window, the centroids of detected cell islands are shown with red marks “*”. Click the “DELETE” button on the MATLAB GUI and select unwanted red marks on the figure window to delete unwanted spots. Once blue circles surround the unwanted red marks, the coordinates of deleted spots are removed. After deletion of all unwanted spots, re-click the “DELETE” button to stop the delete function. Click on the figure window to add red marks directly. All existing and added coordinates of spots are saved.
**2.4.5** After editing, click the “QUIT” button on the MATLAB GUI. The MATLAB program stops, and MATLAB’s interface is minimized by AMFIP. The final coordinates are saved in a text-format list, named “data.txt”. μManager uses this list for the following steps during the experiment.
**2.4.6** The coordinates mentioned in step 2.4.5 are used to guide 40× bright-field acquisition. For each FOV, the MATLAB program is activated by AMFIP and begins to analyze the acquired 40× image for cell islands’ drifting.
**2.4.7** If the cell island’s position has drifted in a 40× image, AMFIP will need new coordinates to replace the current coordinates. First, the MATLAB program informs AMFIP and AMFIP manipulates the XY motor-stage back to the original FOV through μManager. Second, AMFIP controls the camera to take a 10× bright-field image. Third, AMFIP activates the MATLAB program to analyze this image to form a new list of coordinates.

In this analysis, our MATLAB program calculates the distance between the position with drifting and each position in the new list respectively. It will compare all values to get a minimum distance value. If the minimum distance is smaller than 90 μm (the maximum distance of adjacent spots on square lattice pattern. **Fig.4**), this minimum distance value will be accepted, and its corresponding coordinates will replace the coordinates of the position with drifting. AMFIP uses the edited coordinates for 40× bright-field image acquisition and carries on the following operations automatically. If the minimum distance is larger than 90 μm, which means that the cell island in this position has dissolved and disappeared, AMFIP will skip this position and carry on the operation on the next position automatically.

### 3. Cell line generation

Generation of endogenously tagged mNeonGreen2_1-10/11_ cell lines was performed in the human bronchial epithelial cell line (Beas2B) as previously described.[12] Briefly, the DNA sequence coding the 11^th^ strand of fluorescence protein mNeonGreen2 is inserted into the gene of interest (i.e., YAP genomic locus) through the CRISPR-Cas9 gene-editing system, and it complements the 1-10^th^ strand of mNeonGreen2 to emit fluorescence and cells with the tagged protein of interest can be collected through fluorescence-activated cell sorting. As a result, mNeonGreen2 is tagged to YAP whenever the cell expresses YAP in the context of its native gene regulatory network. The “knock-in” cell lines are ready to be used without additional exogenous transfections and can be stably maintained for generations. Correct integration of mNeonGreen211 was confirmed by genomic sequencing and by reduction in fluorescence upon gene knockdown.

### 4. Cell Lines Maintenance

The Beas2B cell line was maintained in humidified incubators with 5% CO2 at 37 °C. Beas2B and endogenously tagged derivatives were cultured in RPMI-1640 medium supplemented with 10% FBS and penicillin-streptomycin at 100 μg/mL. All cell lines were tested for mycoplasma every 3 months using MycoAlert Mycoplasma Detection Kit (Lonza, Basel, Switzerland). All cells used were <20 passages from thaw.

### 5. Cell Imaging

The following steps show how to achieve a multi-functional and long-term image acquisition using AMFIP to observe traction dynamics and YAP dynamics of the YAP-B2B cell line.

**5.1** Turn on the Nikon A1R confocal microscope system following a specific sequence: LU-N4 laser unit, confocal controller, Ti2-E microscope controller, and Ti2-E inverted microscope.
**5.2** In the Ti2-E inverted microscope, switch to 10× objective and the light-path on the right side for bright-field imaging to identify the cells of interest. Using the 10× magnification, open μManager and move XY motor-stage by joystick to find appropriate FOVs containing both single cells and multiple adjacent cells that grow well on the substrate. For each 10× FOV, switch to 40× objective, adjust XY motor-stage again to have the specified FOVs in the center, and record coordinates of selected FOVs.
**5.3** Input these coordinates and predefined experimental parameters into the GUI of AMFIP. For the experiment described in Results F, 40× objective and 5% of DiaLamp intensity are applied.
**5.4** Launch the Elements platform, open the FITC channel, and switch to the resonant scanner for fast-speed imaging. In the experiment described in Results F, fluorescent images captured in the FITC channel display the YAP dynamics of B2B cells that express YAP: mNeonGreen21-10/11. Slowly adjust the knob of the Z-plane and record the highest and the lowest Z position to form a z-stack that covers the overall z-height of cells that start adhering to the substrate.
**5.5** Open DAPI channel and close FITC channel. In the experiment described in Results F, fluorescent images captured in the DAPI channel present displacement of beads that can be used to calculate traction dynamics. Next, slowly adjust the knob to change the Z-plane and record the highest and the lowest Z position to generate a z-stack that covers the interface between the top surface of the substrate and cell bottom.
**5.6** In the macro editor, to generate z-stack images for both laser channels, write specified commands and input (a) 4 quantitative values collected from previous steps and (b) an appropriate step size of z-plane to generate sufficient numbers of frames for a 3D z-stack image. Next, switch back to galvano scanner for high-resolution imaging. To avoid photobleaching of fluorophore and capture images that have low noise, we set the exposure time to 4 seconds for the above experiments.
**5.7** Complete the rest of the commands in macro to achieve the following functions in sequence:
  a. Close DiaLamp and switch to the light-path on the left side for fluorescent imaging.
  b. Switch to the FITC laser channel and start z-stack image acquisition.
  c. Switch to the DAPI laser channel and start a z-stack image acquisition of beads.
  d. Save the two z-stack images to a specified directory for data analysis.
  e. Switch back to the light-path on the right side and turn on the DiaLamp that allows μManager to take a bright-field image.
**5.8** Back to the GUI of AMFIP. To avoid photobleaching, set the time interval for image acquisition of each FOV to 30 minutes. Next, set the total duration of the experiment to 12 hours or above. Next, click “Enter” to start the imaging process.
**5.9** For each time interval (i.e., 30 minutes for the experiment described here), AMFIP automatically executes the following operations in sequence:
  a. Move XY motor-stage to each pre-selected FOV.
  b. Take and save separate z-stack images for FITC and DAPI channels through Elements.
  c. Capture and save a bright-field image through μManager.
  d. After all imaging processes are completed in one FOV, AMFIP automatically instructs XY motor-stage to move to the next FOV and repeat the previous operations.

Each imaging condition of multi-channel images is listed below:

a. Bright-field image: magnification: 40×; DiaLamp intensity: 5%; exposure time: 1s.
b. Z-stack image of DAPI channel: magnification: 40×; laser intensity: 30%; gain of photomultiplier tube: 125; exposure time: 4s; step size: 5 μm; the range of Z-plane: 10 μm.
c. Z-stack image of FITC channel: magnification: 40×; laser intensity: 50%; gain of photomultiplier tube: 70; exposure time: 4s; step size: 2 μm; the range of Z-plane: 30~40 μm.

### 6. Images Processing and Analysis

The ratio of fluorescence intensity between nucleus and cytoplasm (N/C ratio) is widely used in live-cell experiments for the analysis of protein dynamics and cell functions. To examine the relationship between YAP N/C ratio and cell-area/nucleus-area ratio, we analyzed time-lapsed images of spreading cells and non-spreading cells (**Figs. 5C1-5D3**). The following steps show how to use Fiji ImageJ to process and analyze the acquired confocal and BF images.[32]

**6.1** Launch Fiji ImageJ, open all BF images from one FOV, and concatenate them into a stack. Next, open the confocal z-stack image of YAP from the same FOV.
**6.2** Scale down and fit the size of the BF image stack to the size of the confocal image. Go to “Image->Overlay->Add Image”, select the BF image as “image to add”. Next, Set the value of opacity to 50 and click “OK”. This process allows us to overlap the BF image on the confocal image.
**6.3** Locate the cell being examined and align the cell boundary in the BF image with similar shaped YAP fluorescence in the confocal image.
**6.4** Go to “Analyze->Set Measurement”, and check functions: “Area”, “Mean grey value”, and “Integrated density”. For the experiment described in Results F, “Area” measures the area of the selected region of interest (ROI) from the image being processed. “Mean grey value” presents the relative YAP density of the ROI in this experiment. “Integrated density” displays the relative YAP intensity of the ROI which is the product of the data from “Area” and “Mean grey value”.
**6.5** Choose “Freehand selection” on the main interface of ImageJ, first carefully select the ROI of the nucleus of the examined cell and click “Analyze->Measure”. A “Results” window pops up. Second, select a new ROI of the cell boundary of the same cell and redo “Analyze->Measure”.
**6.6** Repeat step 6.5 for every frame of the confocal image stack. Next, copy the data from the “Results” window and paste it into an excel file for data analysis.
**6.7** To determine the YAP density in the cytoplasm, first calculate the difference between the nucleus area and the cell-body area which represents the cytoplasm area. Second, calculate the difference between the integrated density of the cell-body and the nucleus. This value represents the relative intensity of YAP in the cytoplasm. Third, calculate the YAP density by dividing the YAP intensity in the cytoplasm by the cytoplasm area.
**6.8** To calculate the YAP N/C ratio, divide the YAP density of the nucleus by the YAP density of the cytoplasm.
**6.9** Organize the data groups into three columns: YAP N/C ratio, cell area, and nucleus area. Each row of a column represents different time points for imaging. To minimize the influence of photobleaching and the discrepancy of multiple cells, we apply normalization to the data by dividing all the values of each column with respect to the first value of that column, which is the data processed based on the image taken at the beginning of the experiment with no photobleaching.
**6.10** Repeat step 1 to step 9 for all cells being studied. Next, form and analyze multiple line charts based on the processed data (**Figs. 5C1-5D3**).

### 7. Statistics Analysis

We applied a student’s t-test to evaluate the statistical significance of the data (shown in **Figs. 5C & 5D**). To execute this t-test, we input two different data groups for comparison. The calculation generates a p-value that indicates whether there is a significant difference between the two data groups (i.e., p-value > 0.05 means not significant; p-value ≦ 0.05 means significant; “*” means p-value ≦ 0.05, “**” means ≦ 0.01, “***” means ≦ 0.001, “****” means ≦ 0.0001). For both spreading and non-spreading cells, we formed four data groups: normalized YAP N/C ratio, normalized cell area, normalized nucleus area, and normalized nucleus-area/cell-area ratio. We compared the data of each group after 8.5 hours and 9 hours with the data of the same group after 0 hours and 0.5 hours. The calculated p-value indicated whether there is a significant statistical difference between the data at the beginning of cell spreading and the data at the end of cell spreading. Next, we compared the YAP N/C ratio of spreading cells after 8.5 and 9 hours with the YAP N/C ratio of non-spreading cells at the same time points through t-test.

## Acknowledgements

This project is financially supported by Cancer Pilot Award from UF Health Cancer Center (awarded to Drs. Xin Tang and Dietmar Siemann) and the start-up package of Dr. Xin Tang. We sincerely appreciate the intellectual discussions with and the technical supports from Dr. Jonathan Licht (UFHCC), Dr. Rolf Renne (UFHCC), Dr. Hitomi Yamaguchi Greenslet (MAE, UF), Dr. David Hahn (University of Arizona), Dr. Jack Judy (ECE, UF), Dr. Weihong Wang (Oracle Corporation), Dr. Youhua Tan (Hong Kong Polytechnic University), and Support Team of Nikon (Drs. Jose Serrano-Velez, Larry Kordon, and Jon Ekman). We are deeply grateful for the generous and effective supports from all members of Tang’s, Siemann’s, and Guan’s research laboratories and all staff members of the MAE Department.

